# CA inhibitor 1 treatment has only transient effects on estrous cyclicity in mice

**DOI:** 10.64898/2026.01.12.699062

**Authors:** Laura L Giacometti, Donna Mae Dalere, Bupe Lwamba, Samuel Goldberg, Kaitlyn Feliciano, Jacqueline M Barker

**Affiliations:** Department of Pharmacology and Physiology, Drexel University College of Medicine, Philadelphia, PA, USA

**Keywords:** Pre-exposure prophylaxis, estrous cycle, mPOA, astrocytes

## Abstract

Approximately 45% of new HIV infections worldwide occur in women and girls and adherence to pre-exposure prophylaxis (PrEP) to prevent HIV infection is limited. Long-acting PrEP such as lenacapavir may improve adherence. One barrier to PrEP usage in women and girls is impact on the menstrual cycle, though impacts of long-acting PrEP are unclear. To model the effect of lenacapavir administration on cyclicity, adult female C57Bl/6J mice were injected with CA inhibitor 1 and assessed for estrous cyclicity. Ca inhibitor 1 transiently delayed the start of the next cycle. However, Ca inhibitor 1 was associated with a higher proportion of normal length estrous cycles. Ca inhibitor 1 did not impact astrocyte immunoreactivity or chronic cellular activity in the medial preoptic area (mPOA), a major regulator of the estrous cycle. These findings suggest that lenacapavir may be a viable PrEP alternative for those concerned with PrEP impacts on menstrual cycle.

## Introduction

Approximately 41 million people in the world are living with HIV, over half of whom are women and girls. Pre-exposure prophylaxis (PrEP) is highly effective at preventing HIV infection. However, PrEP adherence has historically been limited in women. Lenacapavir, a long-acting multi-stage capsid inhibitor, was demonstrated to yield high safety and efficacy in HIV prevention in women following twice yearly injections (Bekker et al. 2024). Women and girls indicate preference for long-acting PrEP formulations (Little et al. 2024), which may increase PrEP adherence. Women and girls also indicate a preference for PrEP combined with hormonal contraceptives (Little et al. 2024). Further, half of women indicated that PrEP effects on the menstrual cycle would impact choice of PrEP product (Minnis et al. 2019) and PrEP is often used in discordant partners when trying to conceive (Heffron et al. 2018). Thus, it is important to determine any independent effects of lenacapavir on hormonal cyclicity.

Estrous cyclicity is regulated by the medial preoptic area (mPOA) of the hypothalamus (Leedy 1984). The mPOA contains gonadotropin releasing hormone (GnRH) neurons which regulate release of luteinizing hormone and follicle stimulating hormone from the pituitary gland. This in turn controls release of estradiol and progesterone from the ovaries which feedback on the mPOA to regulate its activity.

Neuronal activity in the mPOA is tightly regulated to time surges of luteinizing hormone and follicle stimulating hormone. This coordinated neuronal activity within the mPOA is highly regulated by astrocytes (Sinchak et al. 2020). The astrocyte cytoskeletal marker, glial fibrillary acidic protein (GFAP) is regulated across the estrous cycle in the hypothalamus in rodents (Cashion et al. 2003). Thus, lenacapavir effects on mPOA neuronal activity or astrocyte cytoskeleton may reflect perturbations in regulation of estrous cycle.

To determine if long-acting PrEP formulations impact cyclicity and neural correlates in a rodent model, the effects of the mouse analog of lenacapavir, CA inhibitor 1, on estrous cycle in the female mouse was investigated. Findings indicate that while CA inhibitor 1 transiently increased the latency to the next estrous cycle after administration, CA inhibitor 1 increased the percentage of normal length estrous cycles overall. Further, CA inhibitor 1 did not impact ovarian weight. Immunohistochemical analysis of the mPOA revealed no effect of CA inhibitor 1 on astrocyte reactivity or chronic cellular activity. These findings suggest that lenacapavir may be a viable alternative for women concerned about interactions of PrEP with the menstrual cycle.

## Materials and Methods

### Subjects

Adult female C57Bl/6J (9 weeks, n = 15) mice were obtained from Jackson Laboratories and group housed in same sex cages at Drexel University College of Medicine under standard 12h:12h light conditions and had *ad libitum* access to food and water. All procedures were approved by the Institutional Animal Use and Care Committee at Drexel University.

### CA inhibitor 1 administration

Adult female mice (n = 15) were subcutaneously injected with GS-6207 analog, CA inhibitor 1 (15mg/kg, MedChemExpress, Monmouth Junction NJ, HY-124594), a long-acting HIV capsid inhibitor, or 5% DMSO in sesame oil. This dose was based on a previous study which showed high antiviral efficacy in a humanized mouse model (Yant et al. 2019).

### Vaginal Cytology

Mice were monitored for phase of the estrous cycle using vaginal cytology following treatment with CA inhibitor 1 or vehicle. Vaginal lavage was performed daily at approximately 9:00 (2 hours into the light cycle) using sterile saline to determine the effect of CA inhibitor 1 on estrous cyclicity. The diestrus phase was characterized by predominantly small, round leukocytes; proestrus by round, nucleated epithelial cells; estrus phase by large, irregular, non-nucleated cornified cells; metestrus by a mix of leukocytes, cornified, and nucleated epithelial cells (McLean et al. 2012). Percent time in each estrous phase and average length of the estrous cycle were determined for each mouse.

### Tissue Processing and Immunofluorescence

Ovaries were collected prior to perfusion at 8 weeks post-injection. Brains were collected following perfusion with 4% paraformaldehyde. Following cryoprotection in 30% sucrose, 40 µm coronal sections of the mPOA were obtained using a cryostat. To assess astrocyte reactivity and neuronal activity, sections were blocked using 5% normal donkey serum, incubated in anti-GFAP primary (1:10,000, Sigma-Aldrich, St. Louis MO, G9269) or anti-Δ fos B (1:5000, Cell Signaling Technologies, Danvers MA, 14695) overnight, followed by incubation in biotinylated donkey anti-rabbit secondary (1:500, Jackson ImmunoResearch, 715-085-152) for 30 min. Sections were mounted on plus slides and coverslipped with DPX mounting medium (Electron Microscopy Services).

To analyze GFAP immunoreactivity, 20X images of the mPOA were taken and stitched together using Microsoft Image Composite Editor. The percent area of immunoreactivity was measured using a thresholding method in FIJI. To analyze ΔfosB immunoreactivity, 10X images of the mPOA were taken and stitched together using Microsoft Image Composite Editor. Three composite images were manually outline using FIJI and ΔfosB-immunoreactivity cells were counted using an automate counting method. Counts per area (mm^2^) were average across the 3 sections of the mPOA (AP +0.5 mm, +0.14 mm, -0.22 mm relative to bregma) per mouse. Due to technical issues, 2 mice were excluded from the delta fos B immunohistochemistry and 1 mouse from the GFAP immunohistochemistry analysis.

### Experimental design and statistical analysis

Latency to the first proestrus, average cycle length, ovarian weight, GFAP and ΔfosB immunoreactivity were analyzed using unpaired two-tailed t tests. Average cycle length across the 5 cycles was analyzed in GraphPad using repeated measures 2-way ANOVA for comparing vehicle and CA inhibitor 1-treated mice. Distribution of short, normal length, and long estrous cycles were analyzed by a chi square goodness of fit test.

## Results

### Effects of PrEP on estrous cycle

To determine if long-acting PrEP disrupted estrous cycle, C57Bl/6J female mice received a single injection of CA inhibitor 1 (15 mg/kg) and were monitored for estrous cycle for 5 cycles. Mice receiving CA inhibitor 1 exhibited a significantly longer time to the next proestrus phase following injection [t(13) = 2.227, p = 0.0443] (**Figure 1A**). A mixed effects analysis of average estrous cycle length across the 5 cycles revealed no effect of treatment [F(1, 13) = 0.1029, p = 0.75357], cycle [F(3.039, 39.51) = 2.198, p = 0.1028; Greenhouse-Geisser corrected], and no interaction [F(4,52) = 0.7579, p = 0.5574] (**Figure 1B**). A t test also revealed no effect of CA inhibitor 1 treatment on average cycle length [t(13) = 0.3207, p = 0.5034] (**Figure 1C**). A chi-square goodness of fit test revealed a significant difference in the proportion of normal, short, and long length estrous cycles in mice receiving CA inhibitor 1 compared to vehicle-treated mice [Χ^2^ = 17.2704, p = 0.0083] (**Figure 1D**), consistent with an increased proportion cycles of normal length in mice receiving CA inhibitor 1. Finally, ovarian weights did not differ between CA inhibitor 1 and vehicle-treated mice [t(13) = 0.7215, p = 0.4834] (**Figure 1E**).

**Figure 1.**
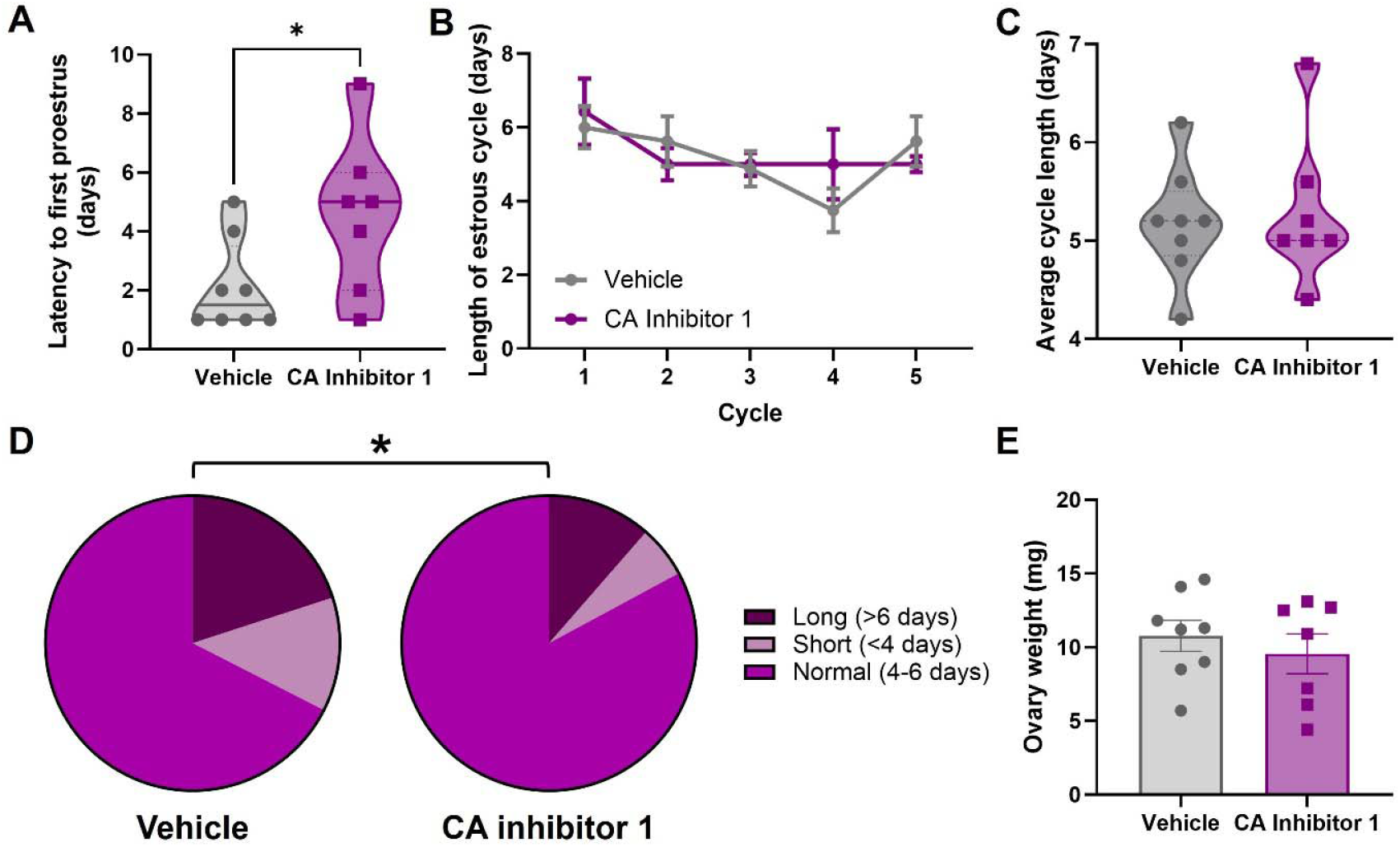
Effects of CA inhibitor 1 administration of on estrous cycle and ovarian weight. (A) CA inhibitor 1 administration resulted in a significant delay in the start of the next estrous cycle. (B) CA inhibitor 1 administration did not impact estrous cycle length across the 5 cycles. (C) CA inhibitor 1 did not impact average estrous cycle length. (D) CA inhibitor 1 significantly increased the proportion of normal length cycles compared to vehicle-treated mice. (E) CA inhibitor 1 did not impact ovarian weight.

### Effects of PrEP on GFAP expression

To determine if CA inhibitor 1 impacted astrocyte reactivity, brain tissue from CA inhibitor 1-treated mice underwent immunohistochemical analysis and quantification of GFAP immunoreactivity in the mPOA (**Figure 2A**). CA inhibitor 1 treatment did not impact GFAP immunoreactivity in the mPOA [t(12) = 0.01862, p = 0.9854] (**Figure 2B-2D**).

**Figure 2.**
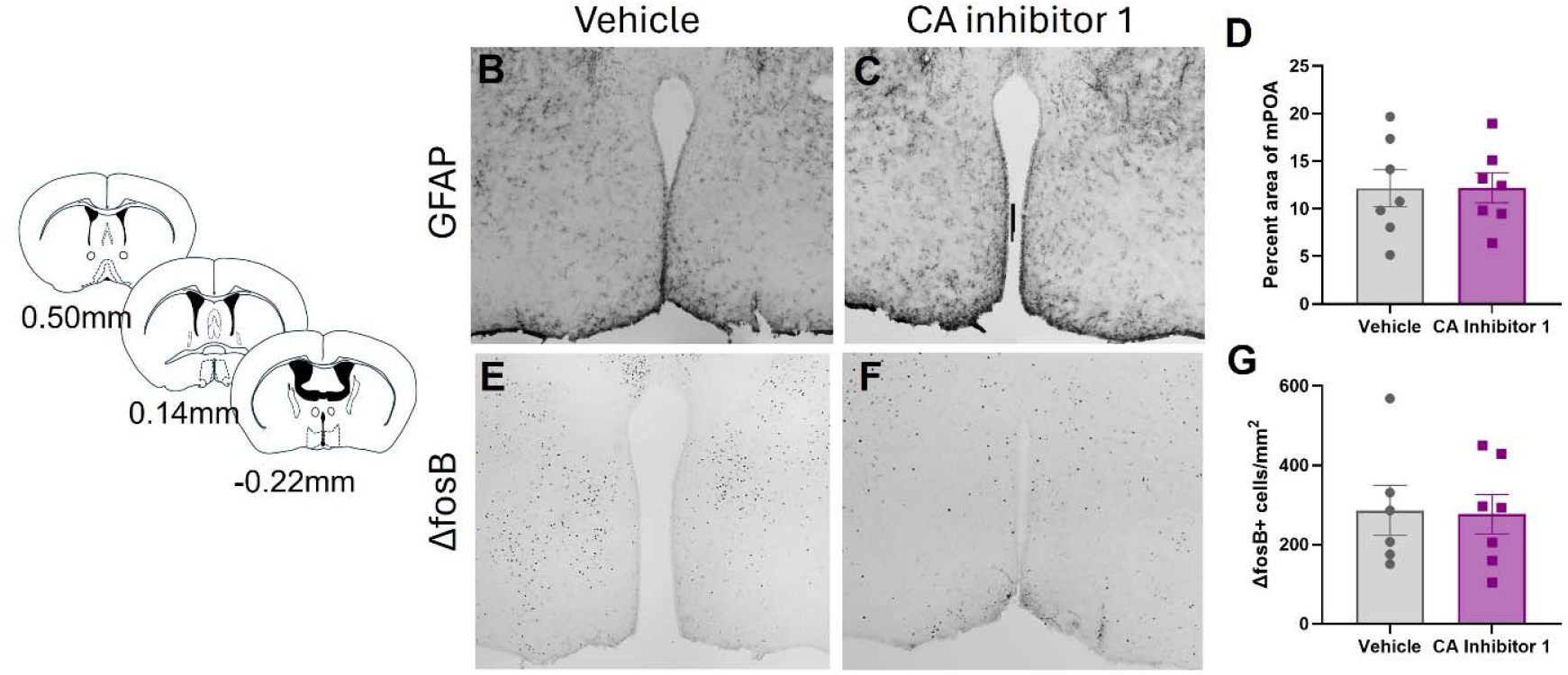
Effects of CA inhibitor 1 on astrocyte reactivity and cellular activity in the mPOA. (A) Schematics of location of mPOA. (B) Representative image of GFAP immunoreactivity in the mPOA of vehicle-treated mice. (C) Representative image of GFAP immunoreactivity in the mPOA of Ca inhibitor 1-treated mice. (D) CA inhibitor 1 did not impact GFAP immunoreactivity in the mPOA. (E) Representative image of ΔfosB immunoreactivity in the mPOA of vehicle-treated mice. (F) Representative image of ΔfosB immunoreactivity in the mPOA of CA inhibitor 1-treated mice. (G) CA inhibitor 1 did not impact ΔfosB immunoreactivity in the mPOA.

### Effects of PrEP on ΔfosB expression

To determine if CA inhibitor 1 impacted neuronal activity, brain tissue from CA inhibitor 1-treated mice underwent immunohistochemical analysis and quantification of ΔfosB immunoreactivity in the mPOA (**Figure 2A**). CA inhibitor 1 treatment did not impact ΔfosB immunoreactivity in the mPOA [t(11) = 0.1224, p = 0.9048] (**Figure 2E-2G**).

## Discussion

Our findings demonstrated that CA inhibitor 1 did not have enduring effects on estrous cycle and, in fact, treatment with CA inhibitor 1 reduced variability in estrous cycle length. Further, CA inhibitor 1 administration was not associated with alterations in astrocyte reactivity or chronic cellular activity in the mPOA in female mice.

Administration of CA inhibitor 1 resulted in a significant delay in the start of the next estrous cycle following injection. While delays in the start of estrous cycle have not been reported on following antiretroviral treatment, acute drug administration has been shown to transiently delay the start of estrous cycles (Kostellow et al. 1980) and repeated drug administration has been shown to result in irregular estrous cycles (King et al. 1993), though this may be dose-dependent (Booze et al. 1999). Following administration of efavirenz and efavirenz combined with tenofovir and lamivudine, prolongation of the estrous cycle in rats has previously been reported (Ohihoin 2018). However, enduring effects of PrEP on ovarian cycle have not been reported. Following CA inhibitor 1 administration, cycle returned to normal. Further, in contrast to what has been observed in rats receiving various combinations of antiretrovirals exhibiting an increase in ovarian weight (Awodele et al. 2018), CA inhibitor 1 treatment did not alter ovarian weights.

CA inhibitor 1 did not impact astrocyte reactivity or ΔfosB immunoreactivity in the mPOA. This suggests that the transient disruption of the estrous cycle by CA inhibitor 1 did not result in long-lasting changes in mPOA activity. However, as the brain tissue was collected after estrous cyclicity had returned to normal, it remains to be determined whether mPOA activity may be transiently altered immediately following CA inhibitor 1 administration. Even transient disruptions in mPOA function may create vulnerabilities to other disruptions such as drug or stress exposure, which independently may impact estrous cycle (King et al. 1993), and may result in more long-lasting changes in estrous cyclicity. As drug use is a major risk factor for HIV infection, future studies should investigate the impact of PrEP on estrous cyclicity following chronic drug use.

One caveat to this study is that mice received a single dose of CA inhibitor 1. While this may more closely resemble the timing of long-acting formulations in humans taking PrEP, which are administered every 2-6 months, there are differences in mouse and human metabolism. Thus, future studies should investigate the long-term effects of chronic dosing paradigms. Another caveat is that brain tissue was not collected in an estrous phase-dependent manner. Previous studies have suggested that GFAP immunoreactivity in the mPOA may be modulated across the estrous cycle (Cashion et al., 2003). While, to our knowledge, ΔfosB immunoreactivity in the mPOA has not been investigated across the estrous cycle, ΔfosB expression is modulated by estrous phase in other brain regions, including the suprachiasmatic nucleus (Shiba et al. 2025). Thus, it is possible that interactions of estrous phase and CA inhibitor 1 administration on immunohistochemical outcomes may have been obscured.

Together, these data suggest that CA inhibitor 1 does not have long-lasting effects on estrous cycle or chronic activity and astrocyte reactivity in a key neural substrate regulating cyclicity, the mPOA. As PrEP adherence is low in women and one of the major factors that mediate PrEP formulation preference is impact on the menstrual cycle, these preclinical findings suggest that lenacapavir may be a promising choice to increase use in women at risk of HIV infection.

## References

Awodele O, Popoola TD, Idowu O, et al (2018) Investigations into the Risk of Reproductive Toxicity Following Exposure to Highly Active Anti-Retroviral Drugs in Rodents. Tokai J Exp Clin Med 43:54–63

Bekker L-G, Das M, Karim QA, et al (2024) Twice-Yearly Lenacapavir or Daily F/TAF for HIV Prevention in Cisgender Women. New England Journal of Medicine 391:1179–1192. 10.1056/NEJMoa2407001

Booze RM, Wood ML, Welch MA, et al (1999) Estrous Cyclicity and Behavioral Sensitization in Female Rats Following Repeated Intravenous Cocaine Administration. Pharmacology Biochemistry and Behavior 64:605–610. 10.1016/S0091-3057(99)00154-9

Cashion AB, Smith MJ, Wise PM (2003) The Morphometry of Astrocytes in the Rostral Preoptic Area Exhibits a Diurnal Rhythm on Proestrus: Relationship to the Luteinizing Hormone Surge and Effects of Age. Endocrinology 144:274–280. 10.1210/en.2002-220711

Heffron R, Thomson K, Celum C, et al (2018) Fertility intentions, pregnancy, and use of PrEP and ART for safer conception among East African HIV serodiscordant couples. AIDS Behav 22:1758– 1765. 10.1007/s10461-017-1902-7

King TS, Canez MS, Gaskill S, et al (1993) Chronic cocaine disruption of estrous cyclicity in the rat: dose-dependent effects. The Journal of Pharmacology and Experimental Therapeutics 264:29–34. 10.1016/S0022-3565(25)10267-X

Kostellow AB, Ziegler D, Kunar J, et al (1980) Effect of cannabinoids on estrous cycle, ovulation and reproductive capacity of female A/J mice. Pharmacology 21:68–75. 10.1159/000137418

Leedy MG (1984) Effects of small medial preoptic lesions on estrous cycles and receptivity in female rats. Psychoneuroendocrinology 9:189–196. 10.1016/0306-4530(84)90038-6

Little KM, Hanif H, Anderson SM, et al (2024) Preferences for Long-Acting PrEP Products Among Women and Girls: A Quantitative Survey and Discrete Choice Experiment in Eswatini, Kenya, and South Africa. AIDS Behav 28:936–950. 10.1007/s10461-023-04202-0

McLean AC, Valenzuela N, Fai S, Bennett SAL (2012) Performing vaginal lavage, crystal violet staining, and vaginal cytological evaluation for mouse estrous cycle staging identification. Journal of Visualized Experiments. 10.3791/4389

Minnis AM, Browne EN, Boeri M, et al (2019) Young Women’s Stated Preferences for Biomedical HIV Prevention: Results of a Discrete Choice Experiment in Kenya and South Africa. JAIDS Journal of Acquired Immune Deficiency Syndromes 80:394. 10.1097/QAI.0000000000001945

Ohihoin DAG (2018) FIRST LINE ANTI-RETRO VIRAL DRUGS DISRUPTS THE ESTRUS CYCLE OF WISTAR RATS. IJHS 3:37–42. 10.47941/ijhs.238

Shiba A, Hardonk MH, Foppen E, et al (2025) Voluntary Running and Estrous Cycle Modulate ΔFOSB in the Suprachiasmatic Nucleus of the Wistar Rat. J Circadian Rhythms 23:7. 10.5334/jcr.257

Sinchak K, Mohr MA, Micevych PE (2020) Hypothalamic Astrocyte Development and Physiology for Neuroprogesterone Induction of the Luteinizing Hormone Surge. Front Endocrinol 11:. 10.3389/fendo.2020.00420

Yant SR, Mulato A, Hansen D, et al (2019) A highly potent long-acting small-molecule HIV-1 capsid inhibitor with efficacy in a humanized mouse model. Nat Med 25:1377–1384. 10.1038/s41591-019-0560-x

